# Minimized double guide RNA libraries enable scale-limited CRISPR/Cas9 screens

**DOI:** 10.1101/859652

**Authors:** Elin Madli Peets, Luca Crepaldi, Yan Zhou, Felicity Allen, Rasa Elmentaite, Guillaume Noell, Gemma Turner, Vivek Iyer, Leopold Parts

## Abstract

Genetic screens based on CRISPR/Cas technology are a powerful tool for understanding cellular phenotypes. However, the coverage and replicate requirements result in large experiment sizes, which are limiting when samples are scarce, or the protocols are expensive and laborious. Here, we present an approach to reduce the scale of genome-wide perturbation screens up to fivefold without sacrificing performance. To do so, we deliver two randomly paired gRNAs into each cell, and rely on recent advances in gRNA design, as well as availability of gRNA effect measurements, to reduce the number of gRNAs per gene. We designed a human genome-wide library that has effective size of 30,000 constructs, yet targets each gene with three gRNAs. Our minimized double guide RNA library gives similar results to a standard single gRNA one, but using substantially fewer cells. We demonstrate that genome-wide screens can be optimized in a demanding model of induced pluripotent stem cells, reducing reagent cost 70% per replicate compared to conventional approach, while retaining high performance. The screen design and the reduction in scale it provides will enable functional genomics experiments across many possible combinations of environments and genetic backgrounds, as well as in hard to obtain and culture primary cells.

## Introduction

Genome-wide genetic screens are a key source of data for understanding gene function. The CRISPR/Cas system has rapidly developed into a tool of choice for these important experiments (Doench 2018). In a typical pooled CRISPR/Cas9 screen, a Cas9-expressing cell line is transduced with a guide RNA (gRNA) library, and propagated for several weeks. The sequence of gRNA in each cell determines the genomic binding of the Cas9 nuclease by annealing to the complementary DNA. Cas9 then cleaves the DNA, ultimately leading to mutations at the targeted locus due to imperfect repair. Frequencies of gRNAs in the pool can be measured by a sequencing readout, and a loss of representation over the course of the screen corroborated by multiple guides indicates that the targeted gene is required for growth in the tested condition.

Scale remains arguably the main limiting factor for screens. Many cell lines have low transduction efficiency, so it is difficult to obtain ample coverage; primary cells are hard or impossible to expand in culture to the extent required; and perturbations transplanted for *in vivo* evaluation tend to become clonal (Chen et al. 2015). Any screen can also be dissected further by repeating it in a different growth condition, a new genetic background, or an additional accompanying mutation. Even a single screen benefits from size reduction to save on cost and labor. Advances in reducing screen sizes would have broad impact.

Several genome-wide human gRNA libraries are publicly available (Shalem et al. 2014; Wang et al. 2015; Doench et al. 2016; Tzelepis et al. 2016; Hart et al. 2017). Early experiments led to appreciating that not all guide RNAs function equally well, and therefore, most libraries contain between four and six gRNAs per gene, resulting in libraries of up to 120,000 constructs. Since then, hundreds of genome-wide screens have been completed, and there is both data on efficacies of cloned gRNAs (Behan et al. 2019; Meyers et al. 2017), as well as improved models for predicting the quality of a new one (Doench et al. 2016). With the help of modelling systems such as JACKS, it is possible to reduce library size up to 2.5-fold, and limit replicates without compromising on the quality of the readout (Allen et al. 2019; Imkeller et al. 2019). Technology has also been developed to introduce multiple gRNAs into the same cell (Doench et al. 2016; Vidigal and Ventura 2015; Wong et al. 2016; Du et al. 2017; Gasperini et al. 2019). Therefore, the scale limitations of genetic screens can be attacked from multiple fronts.

Here, we present an approach to reduce the size of genome-wide CRISPR/Cas9 screens. First, we establish a strategy to make random pairs of gRNAs and introduce them into cells, thus halving the culture volume needed to measure the effect of each guide in isolation. We then use the abundant data on screens in cancer cell lines to prioritize gRNAs to include in a minimized human genome-wide library that targets each gene with three gRNAs. We demonstrate that the library effectively identifies essential genes, and reduces the required experiment sizes for doing so. Finally, we perform screens in human induced pluripotent stem cells, which are considered difficult and cost-prohibitive for this purpose, demonstrating the utility of our approach.

## Results

### Design, cloning and validation of a double gRNA library

We set out to minimize the number of cells required to complete a successful genome-wide screen. To do so, we developed a construct to introduce two gRNAs into each cell (Figure 1A). First, two oligonucleotide pools, each encoding the same set of gRNA specificity sequences, were used as a template in a fusion PCR reaction to produce a large pool of random pairs. We then cloned this pool into an expression vector between the human U6 promoter and gRNA scaffold, and inserted a second scaffold and murine U6 promoter between the two specificity sequences (Figure S1, Methods). The final product encodes a pair of independent gRNA expression cassettes (“5’ gRNA” and “3’ gRNA”) under the control of distinct promoters.

**Figure 1.**
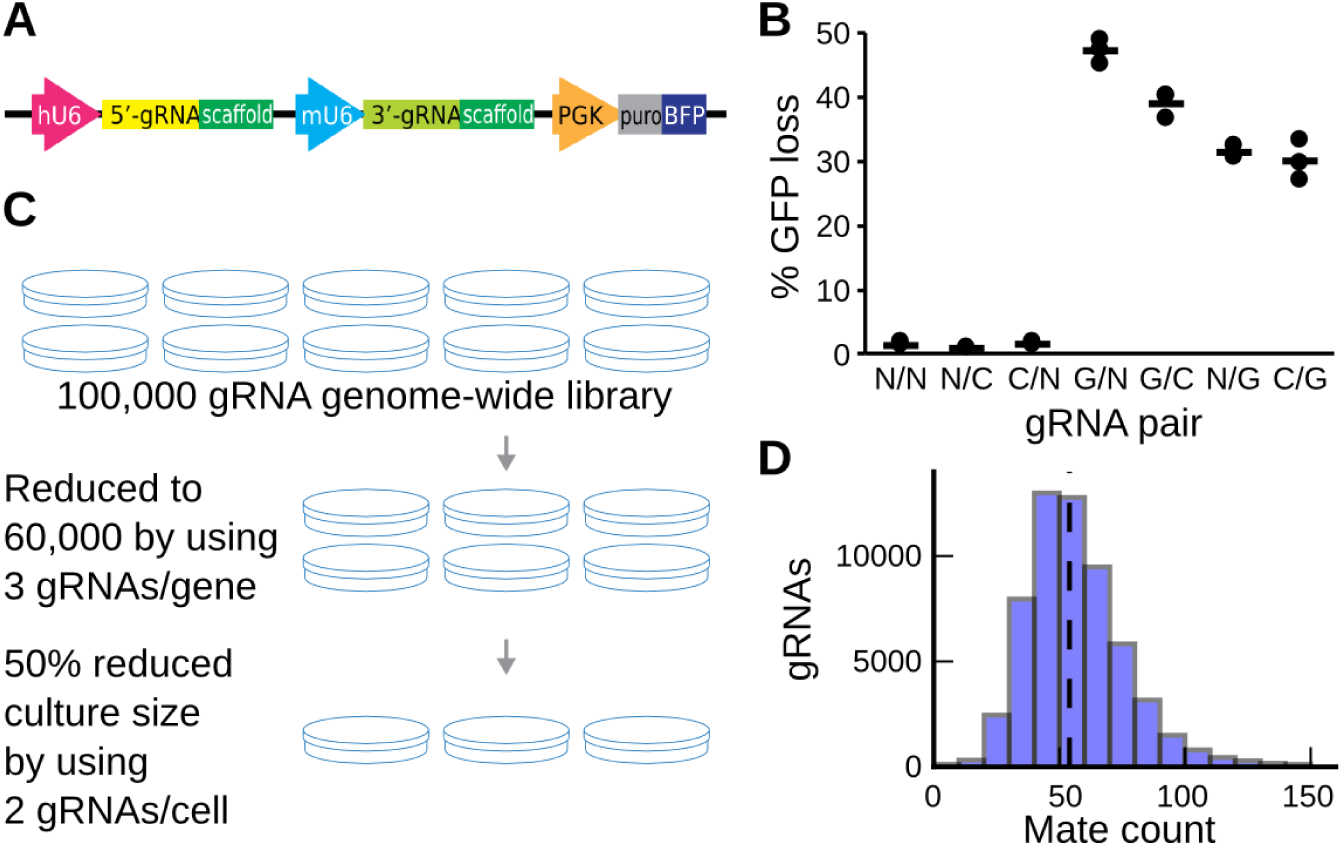
Motivation and design of a double gRNA library. **A.** Design of a construct to express two gRNAs (yellow, green) from the human and mouse U6 promoters (pink, teal), as well as antibiotic resistance and fluorescence markers (grey, blue) from the PGK promoter (orange). **B.** Both 5’ and 3’ gRNAs successfully generate loss of function alleles. Percent of Cas9+/GFP+ K562 cells that have lost GFP fluorescence signal (y-axis) 18 days after infection with pairs of GFP-targeting gRNAs (G), genomic control targeting gRNAs (C) and non-targeting gRNAs (N, x-axis) in triplicate (markers). Lines: averages. **C.** A minimized genome-wide library that uses three gRNAs per gene, and two gRNAs per cell saves 70% of cost per replicate. **D.** Every gRNA has a large number pair mates in the double construct. The number of gRNAs (y-axis) that have a number of different pair mates (x-axis) measured in the plasmid library.

Next, we confirmed that the 5’ and 3’ gRNAs perturb genes with similar efficacy, and are retained upon viral delivery. We transduced K562 cells stably expressing SpCas9 and green fluorescent protein (GFP) with lentiviral vectors expressing pairs of guides targeting GFP or controls, and quantified GFP expression after 18 days by flow cytometry. Both positions were effective at reducing GFP expression, with over 30% of the cells losing GFP signal regardless of the paired partner (Figure 1B). The 5’ gRNA driven from the human U6 promoter was more effective than the 3’ one (43±5% vs. 31±2% average GFP loss), possibly due to higher expression compared to the murine U6 (Roelz et al. 2010). As the double gRNA vector encodes two identical 93bp scaffolds 357bp apart, there is a risk that homologous recombination removes the intervening sequence, including one of the gRNAs (Adamson et al. 2018). We successfully amplified the full double gRNA insert in the genomic DNA of infected K562 cells by using PCR, but could not detect the smaller product that would be generated by recombination events (Figure S2). Thus, between-scaffold recombination occurs at most at negligible levels.

Given both the 5’ and 3’ gRNAs are functional and maintained during cloning and transduction, we proceeded to generate a human genome-wide library targeting 19,259 genes. We have previously shown (Allen et al. 2019) that as few as three gRNAs per gene is sufficient to distinguish gold standard essential genes, as defined by (Hart et al. 2014), from non-essential ones. We therefore cloned three gRNAs per protein-coding gene and non-targeting controls into both positions of the double gRNA plasmid library (Figure S1, Methods). We selected the three gRNAs from the Avana (Doench et al. 2016) and Brunello (Sanson et al. 2018) libraries, and synthesized oligonucleotide pools of gRNA targeting sequences for the two cassettes (Table S1). As we planned to screen in induced pluripotent stem cells (iPSC), we further included a fourth gRNA for 1,986 non-essential genes highly expressed in this cell type. Compared to a conventional library of five gRNAs per gene in a single-gRNA vector, our design reduces the scale of a screen by 70% per replicate (Figure 1C).

There is a concern that unequal representation of gRNAs, or pairing to other gRNAs with large impact will bias gRNA effect estimates, and thereby screen hit identification. To confirm uniform coverage, we sequenced the plasmid library, and in both 3’ and 5’ gRNA positions, found 90% of gRNAs within 50% of median coverage (Figure S3). Notably, the coverages were uncorrelated between the 5’ and 3’ gRNAs (Pearson’s R=-0.02), which will help recover signal for guides that are not well represented in one of the positions. We also tested whether each gRNA is paired to many alternatives, and found that over 94% of gRNAs had at least 30 different partners, and 56 on average (Figure 1D). This high number of partners is expected to not bias individual gRNA effect estimates, provided genetic interactions and positively selected gRNAs are rare (Figure S4).

### Double gRNA library screening quality, scale, and concordance

Provided a functional construct and a well-formed library, we next evaluated its performance in a Cas9-positive chronic myelogenous leukemia cell line K562. We carried out a standard two-week pooled negative selection screen at 134x coverage (Methods), and measured gRNA representation by sequencing the 5’ and 3’ gRNAs independently. The same gRNA produces similar log2-fold changes across replicates (Figure S5), and in the 5’ and 3’ position (R=0.39, Figure 2A), with increased concordance for ones against Hart essential and non-essential genes (Hart et al. 2017). The differences between positions are not explained by experimental noise due to low coverage in the plasmid library, effects of pairing to gRNAs of essential genes, or pairing to non-targeting controls (Figure S6).

**Figure 2.**
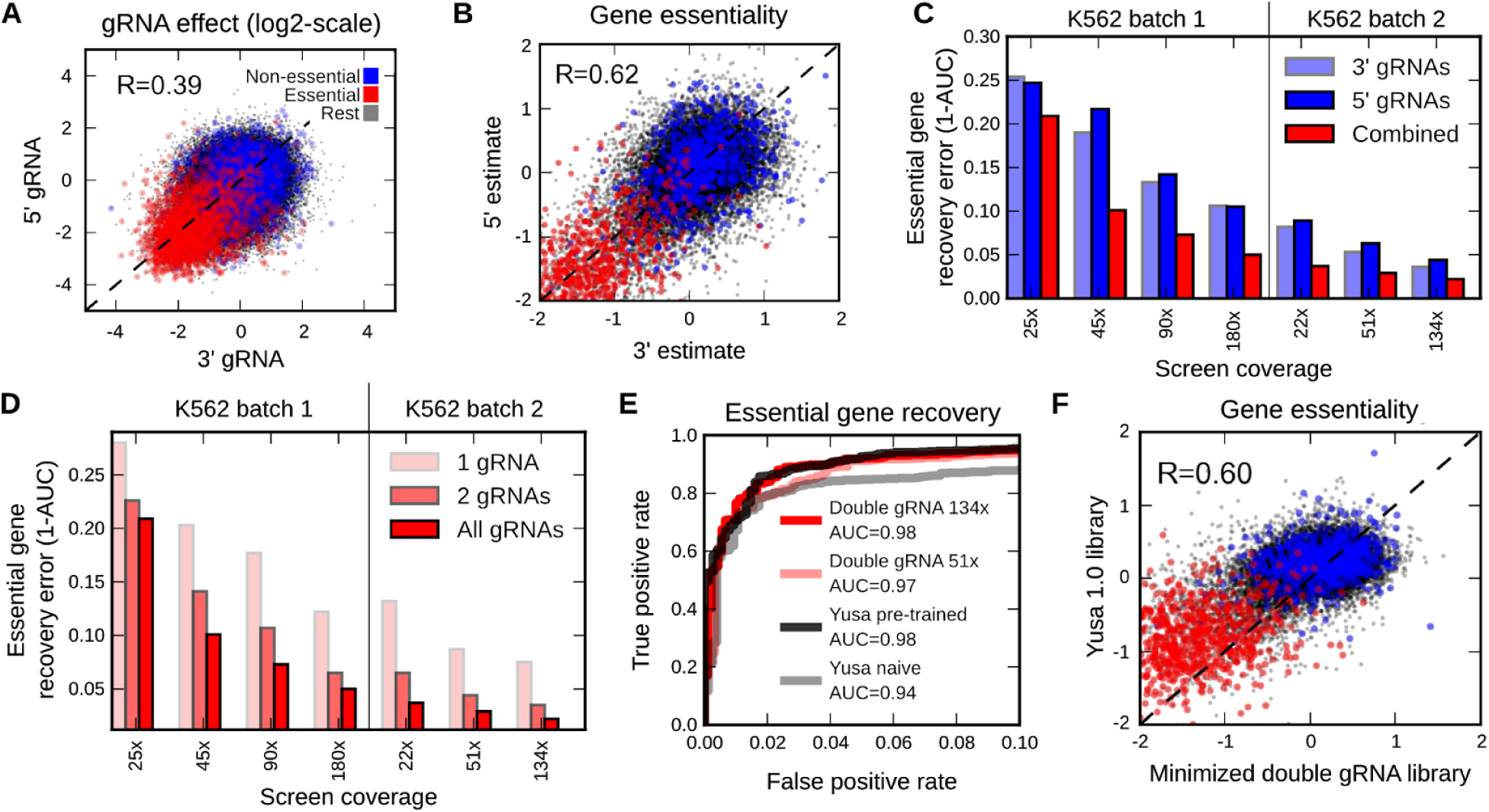
Evaluation of library in K562 cells. All results are from a 134x screen unless indicated otherwise. **A**. gRNA effects correlate between 5’ and 3’ positions. Log2-fold change compared to plasmid control for 3’ gRNAs (x-axis) and 5’ gRNAs (y-axis) targeting essential (red) and non-essential (blue) genes. Dashed line: y=x. **B**. Gene essentiality estimates correlate between 5’ and 3’ positions. JACKS gene essentiality estimated from 3’ gRNAs (x-axis) and 5’ gRNAs (y-axis) targeting essential (red), non-essential (blue), and other (black) genes. Dashed line: y=x. **C**. Combining information from the two gRNA positions improves accuracy on controls. Error in gold standard Hart essential gene classification (y-axis) for different screen coverages (x-axis) using gene essentiality estimates from 3’ gRNAs (light blue), 5’ gRNAs (dark blue), and all gRNAs (red) in two batches of K562 cells (separating vertical line). **D**. Number of gRNAs can be further reduced at the cost of accuracy. Error in essential gene recovery (y-axis) for different screen coverages (x-axis) and increasing number of gRNAs (opacity) for two different batches of K562 cells (vertical separator). **E**. Minimized double gRNA library performs similarly to standard single gRNA one. True positive rate (y-axis) for increasing false positive rate (x-axis, up to 0.10) for distinguishing Hart essential from non-essential genes for double gRNA library at high (dark red) and low coverage (light red) and Yusa 1.0 library analysed naively (grey) or re-using gRNA efficacy estimates from Allen *et al.* (black). **F**. Double gRNA library essentiality estimates are correlated to Yusa library ones. As (B), but for gene essentialities estimated with JACKS from double gRNA library (x-axis) and Yusa 1.0 library (y-axis).

Gene-level essentiality measures are known to be more stable than those of individual gRNAs (Dempster et al. 2019). We used JACKS (Allen et al. 2019) to obtain per-gene estimates that utilize information from all screens performed with this library (Methods). The JACKS gene essentiality measures were indeed better-correlated across replicates (Figure S5) and positions (Pearson’s R=0.62 for all genes, R=0.80 for Hart genes, Figure 2B). Such observed gene effect similarities are consistent with those between screens performed with alternative libraries in different large centres (Dempster et al. 2019). Importantly, the gene effect estimate that combines data from 5’ and 3’ gRNAs improves the ability to distinguish essential from non-essential genes, halving the error (area under the ROC curve (AUC)=0.98 vs 0.96 and 0.96 for combined estimate, 3’ estimate, and 5’ estimate, Figure S7, Table S2). These results demonstrate that both 5’ and 3’ gRNA cassettes contain functional libraries, and that combining information from the two further improves accuracy of separating controls.

The purpose of the downscaled library was to reduce the number of cells in a screen. We therefore tested whether results from a single gRNA screen at a fixed coverage at transduction could be obtained at a lower coverage using the double gRNA library. We ranged the number of successfully transduced cells per construct between 22x and 180x, and evaluated the quality of screen output in two batches of cells. Indeed, we found that using the double gRNA library at a two-fold lower coverage at transduction, and analysing both gRNAs together gives results equivalent to or better than a single guide library (e.g. 1-AUC=0.04 for 22x combined, 0.05 and 0.06 for 51x single from 5’ and 3’ gRNAs, respectively; Figure 2C). On average, using information from two guides per cell compared to a single one resulted in a 49% reduction in error. We also tested whether the number of gRNAs could be reduced further, and re-analysed the data utilising only one or two gRNAs per gene. The second gRNA improved error by 47% on average, and the third one by a further 32% (Figure 2D). The absolute gains were largest for lower coverage screens, highlighting that the total number of cells in which the gene perturbation is assessed is a key parameter for optimization.

The argument to reduce the number of gRNAs was based on statistical considerations (Allen et al. 2019). We next tested whether our new minimized double gRNA library indeed produces gene essentiality estimates that are consistent with established libraries, and can do so with smaller culture volumes. We repeated the screen in K562 cells using the Yusa v1.0 library at a similar 46x coverage per gRNA, and in duplicate. The estimates from both libraries successfully separated essential from non-essential genes (AUC > 0.97, Figure 2E), confirming that there is no reduction of screen output quality from using our selected set of gRNAs, or using the double gRNA construct. The gene essentiality estimates were also correlated (Pearson’s R=0.60, Figure 2F). This level of replication is expected between two different libraries, with average gene effect estimate Pearson’s R of 0.66 across 147 cell lines (Dempster et al. 2019). Altogether, these results justify our approach to reduce the scale of the screens by limiting the number of guides, and increasing the number of gRNAs per cell.

### Double gRNA library enables screens in iPSCs

Screen size is most limiting for cell lines that require expensive media, or have poor transduction efficiency. To demonstrate that the minimized double gRNA library can be used to overcome a challenging screening setup, we tested it in human induced pluripotent stem cells (iPSCs). We generated monoclonal Cas9 positive cells from a donor of the HIPSCI cohort (Kilpinen et al. 2017), and ran a 150x coverage screen (Methods). While there has been some controversy over whether screens are feasible in TP53 wild type cells (Haapaniemi et al. 2019; Brown et al. 2019), especially for cell types with strong response to DNA damage such as iPSCs (Haapaniemi et al. 2018; Ihry et al. 2018), estimates from the screens using our library separate essential from non-essential genes with high accuracy (AUC=0.96, Table S2).

To understand the screen results, we compared the gene essentiality in K562 with that in iPSCs. The gene essentiality estimates are highly correlated (Pearson’s R=0.77 for all genes, R=0.86 for Hart set, Figure 3A), and the core essential genes are shared (intersection/union = 0.76, Figure 3B, Methods), but iPSCs also have additional hits. We used g:Profiler (Raudvere et al. 2019) to test for Gene Ontology categories and Reactome pathways that are differentially essential between the two cell lines (Tables S3, S4). One positively selected set is related to the p53 pathway (“TP53 Network”), which is expected, as it is known that disabling p53-mediated DNA repair signaling increases proliferation in iPSCs (Haapaniemi et al. 2018; Ihry et al. 2018). Several of the genes positively selected in iPSCs upon perturbation (PMAIP1, TP53, CHECK2) were recently individually confirmed in independent studies of stem cell specific essential genes (Mair et al. 2019; Ihry et al. 2019).

**Figure 3.**
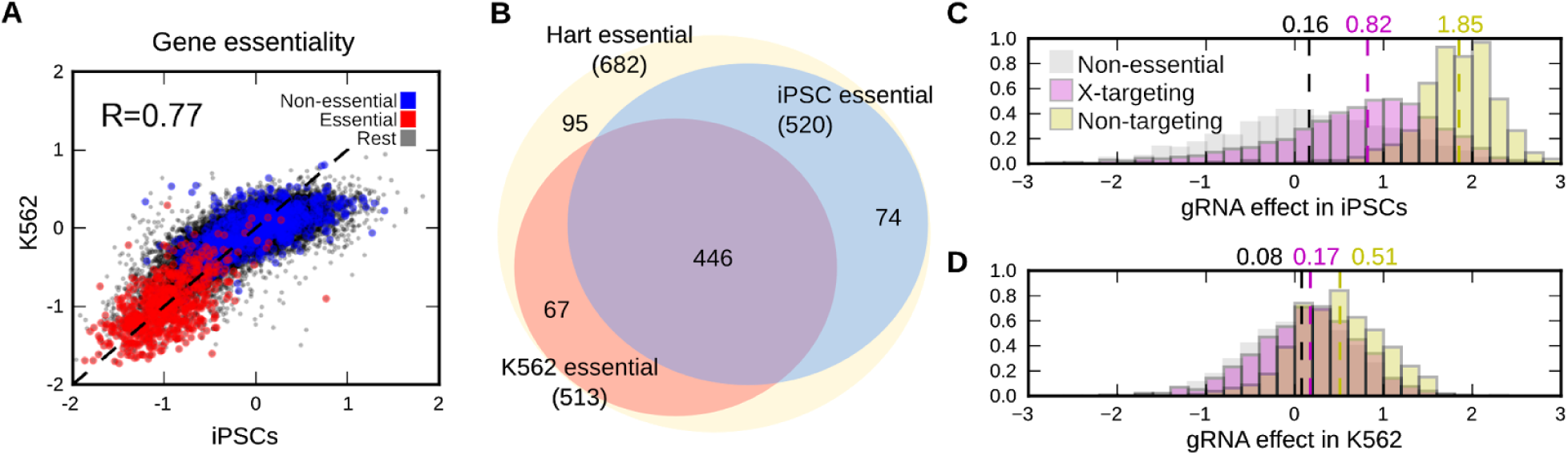
A minimized double gRNA library enables screens in challenging models. **A**. Gene essentialities are correlated between iPSCs and the K562 cell line. Gene essentiality in iPSCs (x-axis) and K562 (y-axis) for essential genes (red), non-essential genes (blue), and the rest (black). Dashed line: y=x; R: Pearson’s correlation. **B**. Essential genes are shared between iPSCs and K562 cells. Venn diagram of Hart essential genes (yellow) overlap with iPSC essential genes (blue), and K562 essential genes (red). iPSC and K562 essential genes not in the Hart set are not displayed. **C**. Additional DNA double strand breaks in a cell retards iPSC growth. Frequency (y-axis) of log2-scale gRNA fold-change in a screen (x-axis) for gRNAs targeting non-essential genes (grey), X-chromosome genes (purple) or not targeting any genes (yellow). Dashed lines and marked values: median effects. **D**. As (C), but for K562 cells.

Cas9-induced DNA double strand breaks are detrimental to growth in screens (Aguirre et al. 2016; Meyers et al. 2017; Gonçalves et al. 2019), and especially so in human iPSCs (Haapaniemi et al. 2018). Therefore, we tested to what extent the additional breaks due to the second gRNA result in unspecific growth defect. We considered gRNAs with decreasing number of target DNA molecules -- two for a non-essential gene targeting gRNA, one for chromosome X-targeting gRNA in the male donor, and zero for non-targeting gRNA. In iPSCs, we observed an average 1.6-fold increase in estimated cell number for gRNAs targeting X, and a 3.2-fold increase in cell number for non-targeting gRNAs (Figure 3C) compared to non-essential gene targeting guides. The effect in K562 cells was substantially smaller (1.5-fold increase in cells with non-targeting gRNAs, Figure 3D) likely reflecting the complex karyotype (Zhou et al. 2019) and DNA repair activity of the K562 cell line. We further performed assays to compare survival and metabolic activity of cells five days post infection for the single gRNA Yusa 1.0 library, and the minimized double gRNA library. At the same infection efficiency of a control line, the double gRNA library had about 25% additional loss of cells, as well as metabolic activity (Figure S8). These results suggest that most of the growth advantage of non-targeting gRNAs is not due to additional cell death, but in shorter arrest to repair DNA before advancing the cell cycle. As the non-targeting controls make up less than 1% of the overall library, their increased representation does not impact the overall screen coverage. In general, a two-gRNA construct reduces scale of cells required at transduction as long as it at most doubles the cell loss at this stage (Figure S9), and halves the cell culture cost for the remainder of the screen.

## Discussion

We described a method to reduce the size of genome-wide CRISPR/Cas screens by evaluating the effects of multiple perturbations in each cell, and reducing the number of gRNAs. We presented a functional two-gRNA vector, demonstrated that the 5’ and 3’ gRNAs are effective in a screen, and that combining information from them further improves hit detection. Given evidence of gRNA efficacy from previous large-scale screens and state of the art prediction tools, we used a data-driven approach to design a new modular human genome-wide library that targets each gene with three gRNAs. This enabled substantial cost and labor savings for screening in the human induced pluripotent stem cells which are difficult and costly compared to cancer cell lines.

Introduction of multiple gRNAs into the same cell has previously been achieved via designed coupling, or increasing the multiplicity of infection. A pair of SpCas9 gRNA expression cassettes was used in genetic interaction screening (Vidigal and Ventura 2015; Wong et al. 2016; Du et al. 2017), and a single construct of four Cas12a crRNAs targeting the same gene was employed to increase the confidence of achieving a complete knock-out (Liu et al. 2019). We opted to pair gRNAs randomly. Under the assumptions of low frequency of positive selection and genetic interactions, which typically hold in a negative selection screen, this reduces confounding from the on- and off-target effects of the limited paired partners for each gRNA. In addition, only single gRNAs need to be synthesized, instead of a quadratic number of combinations required for fully designed pairings, further lowering cost. While higher viral load of a single gRNA library can also increase the number of different gRNAs per cell, it is difficult to control the multiplicity precisely, such that the increased cell death due to transduction, larger number of double strand breaks per cell, and lower chances of a gRNA to end up in a milieu of neutral partners still allow for accurate estimates of its effect. We suggest that this alternative is most useful in settings where infection and double strand break toxicity are not a major concern, and a large majority of gRNAs are expected to not have an effect, such as CRISPR interference or activation screens of regulatory elements.

The benefit of reducing screen scale can be substantial. Using three empirically chosen gRNAs per gene, two gRNAs per cell, two replicates, and more sensitive analysis methods gave a five-fold size reduction over our initial iPSC screens that employed a single gRNA library of five gRNAs screened in triplicate, and analyzed with standard methods. Such reductions can result in a step change in the range of feasible experiments. For example, flow sorting based screens for pathway activity operationalized by antibody staining are a promising, but laborious approach for understanding cell signaling and responses to environment (Brockmann et al. 2017). Sorting for the bottom 1% of the distribution of activity from a pool of cells perturbed with 100,000 gRNAs at 100x coverage with 10,000 cells per second flow rate in triplicate would take about 83 hours. A five-fold reduction (using two replicates) would bring this down to two eight-hour days of sorting, one for each replicate -- a much more practical solution.

The strength of our approach is the flexible combination of control and randomization. The unique PCR handles for each of the three guides and two positions included in the oligonucleotide synthesis phase allowed for different configurations of random guide pairings. This enables customizable experimental design, where the same gRNA sub-pools could be combined depending on the desired tests. In principle, we could further introduce a multi-gRNA expression construct (Liu et al. 2019), include unique molecular identifiers (Michlits et al. 2017; Schmierer et al. 2017), and insert additional expression constructs (Martella et al. 2017) to combine the benefits of the different approaches. Care is needed to ensure equal efficacy and representation of all products in these highly multiplexed settings.

The idea of evaluating the effect of two gRNAs in each cell can be applied to any existing library. However, this requires re-synthesizing the constructs for compatibility with our cloning strategy. The benefits of multiple gRNAs per cell can be reduced if there is excessive cell death, or low enough coverage that the paired mate effects are substantial. We maintained high library coverage in our cloning, with an average of 56 paired partners, and estimated an additional 25% loss of cells due due to the additional DNA breaks, as well as growth retardation for about 1.7 doublings (3.2-fold reduction of cell number compared to controls) in iPSCs. While the cloning coverage can be straightforwardly controlled, the effect of additional DNA breaks should be evaluated to accurately estimate the screen coverage.

Controlled and scalable approaches for perturbation will be key for screens in precious primary samples, and scaling across combinatorial spaces of drugs and genetic backgrounds. Efficient experiments need to strike a balance between increasing signal from each cell by measuring the combined effects of multiple perturbations, and increasing confounding due to resulting biases. Our method prioritizes gRNAs according to evidence from hundreds of screens, introduces two gRNAs into each cell, and uses powerful analysis tools, enabling a range of designs.

## Methods

### Selection of guides and targets

We constructed a new genome-wide library based on the Avana library (Doench et al. 2016) that had ample screening data available to inform gRNA choice (Meyers et al. 2017), and Brunello library (Sanson et al. 2018) that reflects state of the art gRNA efficacy prediction results. We first processed the JACKS results of the Avana dataset (Allen et al. 2019) to identify genes that have essentiality score below −0.7 in at least one cell line. We obtained 5546 essential, and 12095 non-essential genes according to this designation. We ranked the Avana gRNAs according to their median log-fold change across cell lines, and the Brunello gRNAs according to their Rule Set 2 score (Sanson et al. 2018). We then picked the top two data-informed Avana gRNAs for each essential gene, and the top-scoring unused Brunello gRNAs for the third. For non-essential genes, we picked the top two Brunello gRNAs, and one from Avana. If enough gRNAs were not available from one library, top ones from the other were used instead. We further picked two non-overlapping sets of 398 non-targeting gRNAs from the Brunello library, and added one to each of the 3’ and 5’ gRNA constructs. Finally, we added a fourth gRNA for 1986 non-essential genes highly expressed in iPSCs (log2 expression value at least 16) according to (Kilpinen et al. 2017). The selected gRNA targeting sequences were synthesized in an oligonucleotide construct (Figure S10) separately for 5’ and 3’ gRNA libraries. Both 5’ and 3’ libraries thus comprised 13713+13707+13610 gRNAs against non-essential genes, 5546+5544+5496 gRNAs against essential genes, 1986 gRNAs against highly expressed non-essential genes in iPSCs, and 398 non-targeting controls, for a total of 60,000 gRNAs.

### Cloning of double gRNA vectors and library

Double-gRNA expression constructs were cloned starting from pairs of oligonucleotides encoding gRNAs positioned at either 5’ or 3’ cassette. Fusion PCR was employed to join 5’ and 3’ oligonucleotides into a single dsDNA fragment, followed by EcoRV digestion at both ends to expose homology regions required for the subsequent Gibson assembly. Column purification QIAquick PCR Purification Kit (QIAGEN) was employed after each step. The lentiviral backbone vector pKLV2-U6gRNA5(BbsI)-PGKpuro2AmCherry-W (Addgene 67977, (Tzelepis et al. 2016) was linearised with BbsI. EcoRV-digested amplicons encoding gRNA pairs were inserted into the vector by Gibson assembly (NEB Gibson Assembly Master Mix) according to manufacturer’s specifications, and transformed by electroporation (NEB 10-beta Electrocompetent E. coli C3020K). Bacterial cells were cultured overnight and plasmid DNA encoding an intermediate construct was extracted. A dsDNA gBlock (IDT) encoding a scaffold and the mouse U6 promoter (Du et al. 2017), Figure S10) was amplified with KAPA polymerase using primers #545 and #546 (Table S5) and purified with Monarch DNA Cleanup Columns (NEB). Both intermediate construct and amplified gBlock were digested with BbsI (NEB) and ligated using T4 DNA Ligase (NEB), followed by transformation by electroporation.

For the generation of the genome-wide minimized gRNA library, a 143-mer oligo pool encoding 120,000 oligonucleotides (60,000 for each 5’ and 3’ gRNA) was purchased from Twist Bioscience. The pool was composed of several sub-pools to allow for the selective amplification of gRNAs encoded in either the 5’ or 3’ expression cassette in the final product (Table S1). A PCR reaction using KAPA polymerase (Roche), 10 ng template, primers #221 (or subpool-specific)/#270 (Table S5), converted the single-stranded DNA oligos to double-stranded DNA and at the same time amplified position-specific subpools (Figure S1). 5’ and 3’-specific oligos were further amplified by nested PCR using primers #221/#526 and #527/#270 respectively. Finally, fusion PCR was employed to join 5’ and 3’ gRNA sub-pools into a single pool of amplicons composed by high-complexity randomised pairs, followed by EcoRV digestion at both ends to expose homology regions required for the subsequent Gibson assembly. All PCR steps were carried out at low cycle numbers to minimize the risk of amplification bias. Column purification QIAquick PCR Purification Kit (QIAGEN) was employed after each step.

The lentiviral backbone pKLV2-U6gRNA5(BbsI)-PGKpuro2ABFP-W (Addgene 67974, (Tzelepis et al. 2016) was linearised with BbsI. EcoRV-digested amplicons encoding gRNA pairs were inserted into the vector in 3 separate Gibson assembly reactions (NEB Gibson Assembly Master Mix) according to manufacturer’s specifications. Reactions were pooled, column-purified and transformed in 14 electroporations (NEB 10-beta Electrocompetent E. coli C3020K). Bacterial cells were cultured overnight and plasmid DNA encoding an intermediate library was extracted using QIAGEN Plasmid Midi Kit (QIAGEN). A dsDNA gBlock (IDT) encoding a scaffold and the mouse U6 promoter was amplified with KAPA polymerase using primers #545/#546 (Table S5) and purified with Monarch DNA Cleanup Columns (NEB). Both intermediate library and amplified gBlock were digested with BbsI (NEB) and ligated using T4 DNA Ligase (NEB) in three separate reactions. Ligations were pooled, purified with Monarch DNA Cleanup Columns (NEB) and digested again with BbsI to remove any carryover of undigested, intermediate library. Sample was purified and transformed in 20 electroporations to ensure high library complexity (average 170-fold gRNA coverage). Bacterial cells were cultured overnight and plasmid DNA encoding the genome-wide minimized library was extracted using QIAGEN Plasmid Midi Kit (QIAGEN).

### Generation of Cas9 expressing iPSC line and Cas9 activity validation

Cas9 expressing monoclonal iPSC lines were generated using a lentiviral expression vector encoding SpCas9 and a blasticidin resistance (pKLV2-EF1a-Cas9Bsd-W, Addgene, 68343) (Tzelepis et al. 2016). Cas9 expressing lentivirus was packaged using lentiviral production method as described above. Wild-type iPSCs were dissociated with accutase, then 5×10^5 cells were infected with Cas9 expressing lentiviral supernatant in 6-well plate in the presence of 10μM Rock inhibitor Y-27632. After 24h of culturing, the medium was replaced with TeSR-E8 without Rock inhibitor. Blasticidin (TOKU-E, B001) selection was initiated two days after infection with 10μg/ml in TeSR-E8 medium. Thereafter, cells were maintained in Blasticidin medium for 5-7 days. Transduced cells were collected and approximately 600 single cells were subcloned into 6-cm dish pre-coated with 1mg/ml Synthemax™ II-SC Substrate (Corning, 3535) at a concentration of 5 μg/cm^2^. Cells were cultured with TeSR-E8 medium containing 10x CloneR (Stem Cells, 05888) and Blasticidin selection was maintained every other day thereafter. Approximately 8-10 days later, 10-12 single cell derived colonies were manually isolated and transferred into 12-well plates coated with Vitronectin XF (Stem Cell, 07180). Subsequently, the Cas9 expressing monoclonal cells were further expanded for Cas9 activity validation and downstream experiments.

The Cas9 activity of Cas9 expressing monoclonal lines was validated as previously published (Tzelepis et al. 2016) by using lentiviral constructs pKLV2-U6gRNA5(gGFP)-PGKBFP2AGFP-W and pKLV2-U6gRNA5(Empty)-PGKBFP2AGFP-W as a control (Addgene 67980 and 67979 respectively). 5×10^5 Cas9 expressing single cells were infected with either lentiviral supernatant in 6-well plate with 2ml TeSR-E8 in the presence of 10uM Rock inhibitor. Three days post infection, cells were harvested as single cells and analysed by FACS for BFP and GFP expression. Cas9 activity was calculated as percentage of GFP negative vs GFP positive cells in BFP positive cells.

### Cell culture

K562-Cas9 cells were a kind gift by E. De Braekeleer. K562-Cas9-GFP cells were generated by inserting a CMV-GFP expression cassette in the AAVS1 locus. Briefly, K562-Cas9 cells were transiently cotransfected with an expression construct carrying a gRNA targeting the AAVS1 locus (Mali et al. 2013) and a donor plasmid encoding the CMV-GFP expression cassette surrounded by 70 bp homology arms. 14 days later cells were sorted based on GFP expression, single clones were expanded and the correct insertion events were selected by PCR analysis and Sanger sequencing. K562-Cas9 and K562-Cas9-GFP cells were cultured in RPMI supplemented with 10% FCS, 2 mM L-glutamine, 100 U/ml penicillin and 100 mg/ml streptomycin. The cells were treated with 15 µg/ml blasticidin for a week before starting a screen to ensure stable Cas9 expression.

Human iPSCs were cultured on vitronectin-XF-coated plates (Stemcell Technologies) and TeSR-E8 medium (Stemcell Technologies). E8 medium was changed daily throughout expansion and all experiments. All cell lines were cultured at 37 °C, 5% CO2.

### Lentivirus production and determination of lentiviral titer

Supernatants containing lentiviral particles were produced by transient transfection of 293FT cells using Lipofectamine LTX (Invitrogen). 5.4 μg of a lentiviral vector, 5.4 μg of psPax2 (Addgene 12260), 1.2 μg of pMD2.G (Addgene 12259) and 12 μl of PLUS reagent were added to 3 ml of OPTI-MEM and incubated for 5 min at room temperature. 36 μl of the LTX reagent was then added to this mixture and further incubated for 30 min at room temperature. The transfection complex was added to 80%-confluent 293FT cells in a 10-cm dish containing 10 ml of culture medium. After 48 h viral supernatant was harvested and fresh medium was added. After 24h the lentiviral supernatant was collected and mixed with the first supernatant which was then stored at −80 °C. When necessary we prepared larger amounts of lentivirus by scaling up the procedure above.

For gRNA library lentiviral titration, K562 cells were plated into 96-well plate, 5×10^4 cells per well. 8 μg/ml Polybrene (hexadimethrine bromide, Sigma) was added to each well and the cells were transduced with a varying volumes of virus (0 to 20 μl). The cells were then centrifuged at 1000g for 30 minutes at room temperature and resuspended in the same media. After three days of cell culture, cells were harvested for FACS analysis and the level of BFP expression was measured. Virus titer was estimated and scaled up accordingly for subsequent screens.

For gRNA library lentiviral titration on Cas9 expressing iPSCs, Cas9 expressing iPSCs were harvested by accutase (Stem cell, 07920) as single cells. iPSCs (3.6×10^5/well in 6-well plate) were infected with at least five serial dilutions of lentiviral supernatant supplemented with 10uM Rock inhibitor Y-27632 (Stem cell, 72304). Uninfected cells were used as negative control. The transduced cell mixture was cultured in 6-well plates in 1.4ml/well. 24h post transduction, the medium was refreshed with TeSR-E8 without Rock inhibitor. After three days of cell culture the cells were harvested for FACS analysis and the level of BFP expression was measured. Virus titer was estimated and scaled up accordingly for subsequent screens.

### Cell viability assay

To test the effect of lentiviral infection on cell proliferation and viability 5,000/well K562 cells stably expressing Cas9 were plated into 96-well round bottom plates and infected with increasing amounts of lentiviral supernatant. For each volume of lentivirus at least 10 replicates were run. Three days later, cell viability was measured using CellTiter 96® AQueousOne Solution Cell Proliferation Assay (Promega) according to manufacturer’s specifications. On the same day, multiplicity of infection (MOI) was measured by FACS analysis of BFP expression levels. A similar protocol was employed to assess viability in wild type and Cas9-expressing iPSCs, using 7,000 cells/well in 96-well flat bottom plates, and viability and MOI were measured five days after infection. Since in absence of Cas9 the infection with a gRNA library did not trigger the DNA damage stress response, MOI in wild type cells was used as an indirect quantification of lentiviral titer.

### Screening

Cells were infected with the lentiviral construct aiming for MOI of 0.35. Different amounts of cells were infected according to the target coverage. 24h after transduction, 2 μg/ml puromycin was added to K562 cells and the selection was maintained for one week. A small subsample of cells were cultured without puromycin selection to assess the MOI of the screen after 7 days of infection. The screened cells were passaged every 3-4 days for two weeks with at least 200x coverage and at least twice the minimum number of cells required to maintain coverage were spun down and frozen for genomic DNA extraction.

iPSC cells were infected with MOI of 0.3. After 72h, 0.6 μg/ml of puromycin was added and maintained until the end of the screen. Cells were passaged when they reached confluency maintaining at least 200x coverage, and pellets collected for genomic DNA extraction contained at least twice the minimum number of cells required to maintain coverage.

### Genomic DNA preparation and sequencing

For genomic DNA extraction of screen samples, cells were resuspended in 100 mM Tris-HCl, pH 8.0, 5 mM EDTA, 200 mM NaCl, 0.2% SDS and 1 mg/ml Proteinase K and incubated at 55ºC overnight. The following day, samples were treated with 10 µg/ml RNase A for 4h at 37ºC. After adding one volume of 100% isopropanol, genomic DNA was spooled out and washed three times in 70% ethanol. DNA was air dried and resuspended in TE buffer overnight. DNA was quantified using Quant-iT Broad Range kit (Invitrogen). Alternatively, for the PCR analysis of potential recombination events genomic DNA from 1×10^6 cells was extracted using DNeasy Blood & Tissue Kits (Qiagen) following the manufacturer’s instructions, followed by PCR on 500 ng gDNA using primers #1 and #2 (Table S5).

Sequencing libraries were generated through two consecutive PCR reactions. First, gRNA cassettes were amplified separately using Q5 Hot Start High-Fidelity 2X Master Mix (NEB) and primers #1 and #463 for 5’ or #605 and #430 for 3’ cassette. The PCR settings used were 98ºC 30s followed by 98ºC 10sec, 61ºC 15s, 72ºC 40s for 25 cycles, and a final extension at 72ºC 2m. For each sample the amount of genomic DNA to be used was determined by the coverage of the screen and multiple reactions were run when necessary. Reactions for each sample were pooled and purified using QIAquick PCR Purification Kit (QIAGEN). The reactions were eluted into 30 μl of ultra-purified water and quantified with Nanodrop. All the samples were normalised using a concentration of 1 ng/μl. Sequencing adaptors were extended and libraries were indexed in a second PCR using primer #15 and a reverse indexing primer as described before (Tzelepis et al. 2016). PCR products were purified with Agencourt AMPure XP beads, quantified with Quant-iT High Sensitivity kit (Invitrogen), pooled and single-end sequenced using primers #16 and #619 for 5’ and 3’ cassette libraries respectively. For the plasmid library both cassettes were amplified as one amplicon with primers #1 and #2 and sequenced by paired-end sequencing using primers #16 and #436 (Table S5).

### Sequence analysis

We counted sequencing reads that perfectly matched the designed gRNA sequences for each gRNA from the 3’ and 5’ positions of the double gRNA construct. We used JACKS (Allen et al. 2019) with default parameters to infer per-gene and per-sample gene essentiality values for the double gRNA library, using the plasmid counts as controls, and combining timepoints beyond day 10 as replicates. To measure gene essentialities from fewer gRNAs, we alternatively retained only gRNA set 1 for one gRNA, and gRNA sets 1 and 2 for two gRNAs, and otherwise used JACKS using the same way. We computed screen metrics (described below) using the 3’ and 5’ gRNAs as different screen samples, as well as different gRNAs. For Yusa library, we also used the gRNA efficacies pre-calculated in (Allen et al. 2019), and ran JACKS with this additional information.

To evaluate the impact of paired mates, we estimated to summary statistics per gRNA at each of the 5’ and 3’ positions. First, we considered the loss of fitness due to pairing to a deleterious gRNA. We calculated the fitness of each mate pair gRNA as the ratio of frequencies in the target sample and plasmid control, capped at 1, and calculated the expected fitness of the focal gRNA in a sample by averaging fitnesses of all its paired mates, weighted by the number of times they are observed together in the plasmid control. Second, for each gRNA, we calculated the fraction of reads in constructs where the other gRNA is non-targeting in the plasmid library. To simulate mate pairings, we ranged the number of gRNAs against essential genes from 60 to 30,000, and the average mates for each gRNA from 1 to 100, bootstrapping 100 samples, and storing the largest, median, and mean fraction of mates paired to a gRNA against an essential gene.

We used Scikit-learn (Varoquaux et al. 2015) to calculate ROC curves, as well as the corresponding areas under the curve. We calculated other metrics (partial area under the curve, 1-recall at given false positive rate, false positive rate at given true positive rate, delta AUC) as described in (Allen et al. 2019). We used g:Profiler (Raudvere et al. 2019) version e98_eg45_p14_ce5b097 with an ordered query mode on the 250 genes with largest positive and negative differences between iPSC and K562 gene essentiality estimates.

## Code and data availability

Processed files (read counts, log-fold changes, gene essentiality estimates), Jupyter notebooks, and code are available on Figshare: https://figshare.com/projects/Minimized_double_guide_RNA_libraries_enable_scale-limited_CRISPR_Cas9_screens/72296

## Acknowledgements

Author contributions. Performed experiments: EMP, LC, YZ, RE. Developed protocols: EMP, LC, YZ, RE, GT. Analysed data: FA, GN, LP. Supervised work: VI, LP. Wrote paper: EMP, LC, LP. The authors would like to thank Eugene Kwa, and Vitalii Kleshchevnikov for comments on the text. This work was supported by Wellcome (all authors), and Estonian Research Council (IUT34-4).

