## Supplementary Figures and Tables for "Minimized double guide RNA libraries enable scale-limited CRISPR/Cas9 screens"

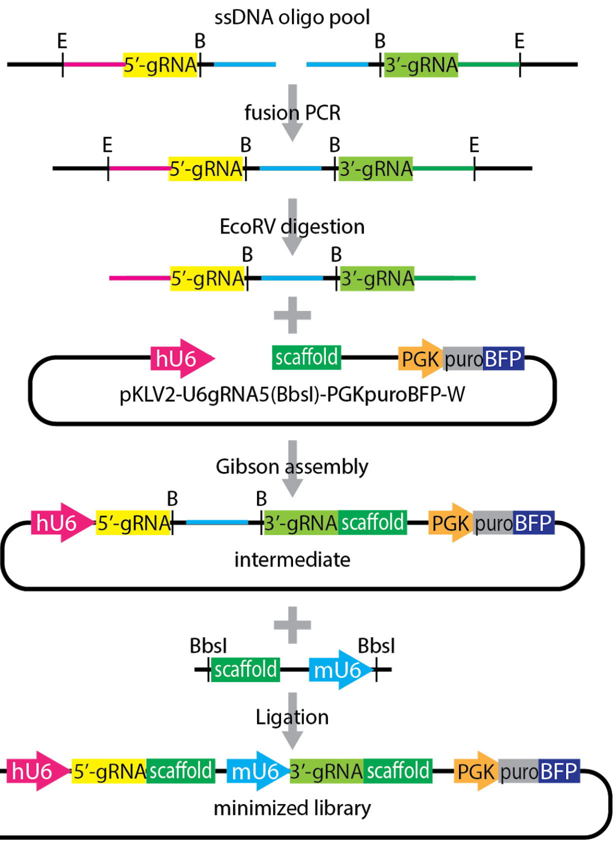


**Figure S1.** Workflow for the cloning of a randomised, double-gRNA library. Blue line - common sequence for fusion PCR; magenta line - common sequence with human U6 promoter; green line - common sequence with gRNA scaffold; E - EcoRV digestion site; B - BbsI digestion site. Other colors and sequences as in Figure 1A. See Methods for details on cloning procedure.

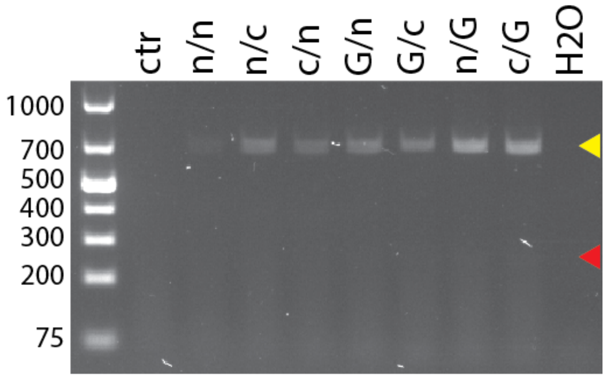


**Figure S2**. Lack of recombination between two gRNA cassettes. K562-Cas9 cells were infected with lentiviral vectors encoding gRNA pairs targeting GFP (G), a control target (c) or non-targeting (n). After 18 days in culture, genomic DNA was extracted and analysed by PCR targeting the full double gRNA insert. The expected 713 bp product corresponding to the intact cassette was detected (yellow arrowhead), but not a 263 bp product (red arrowhead) deriving from the excision of part of the cassette due to homologous recombination.

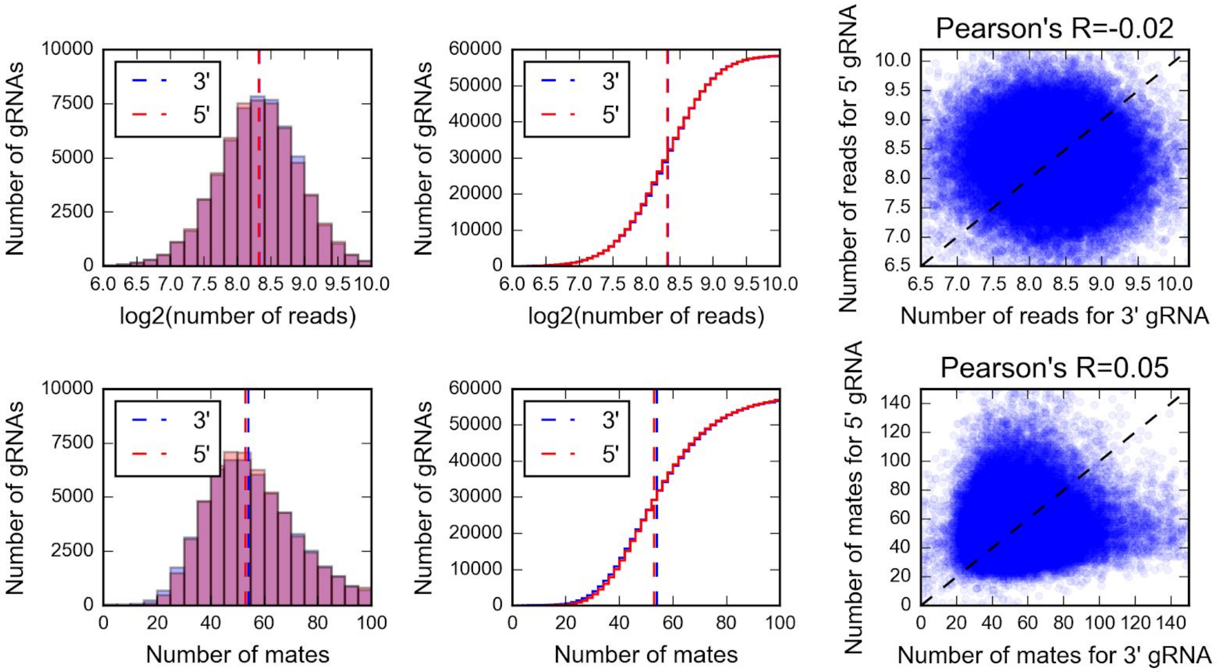


**Figure S3**. gRNA coverages and mate counts in the plasmid library are large, and uncorrelated between the 5' and 3' positions. **A**. gRNAs have high sequencing read coverage from the plasmid library. The number of gRNAs (y-axis) that have a log2-scale coverage (x-axis) measured in the plasmid library for 5' (red) and 3' (blue) gRNAs. Dashed line: median number of reads. **B**. As (A), but cumulative number on the y-axis. **C**. Read counts are uncorrelated between 3' and 5' gRNAs. Log2-scale read coverage for the 3' gRNA (x-axis) and 5' gRNA (y-axis) for each of the 60,000 gRNAs present in the plasmid library (markers). Dashed line: y=x. Pearson's correlation coefficient is marked on the plot. **D-F.** As A-C, but for number of pair mates in the double gRNA construct, counted from the plasmid library sequences.

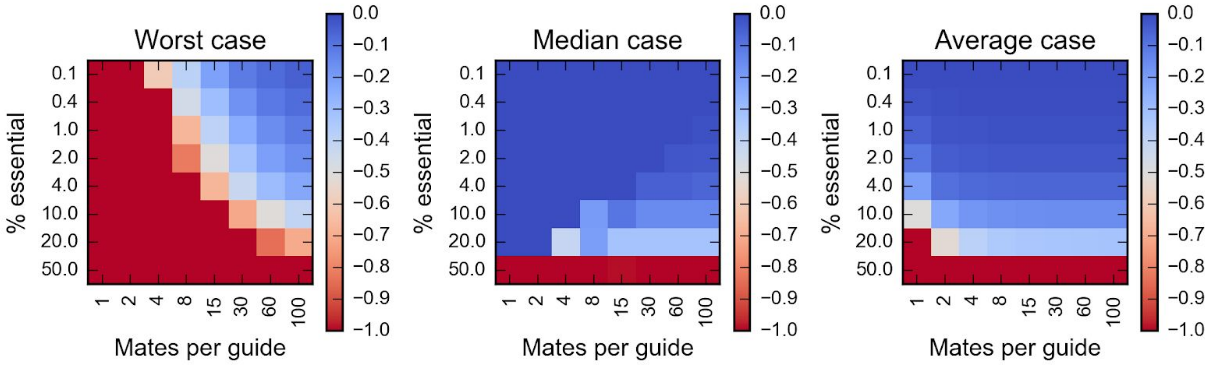


**Figure S4**. Expected bias due to paired gRNAs is small on average. Bias of gRNA effect estimate (log2-fold change compared to control, color) due to pairing to gRNAs targeting essential genes depending on the percent of essential genes (y-axis) and the number of random mates for the gRNA (x-axis). Left panel - worst case (largest bias out of 1000 bootstrap samples); medium panel - median case (median bias out of 1000 bootstrap samples); right panel - average case (average bias out of 1000 bootstrap samples).

A.

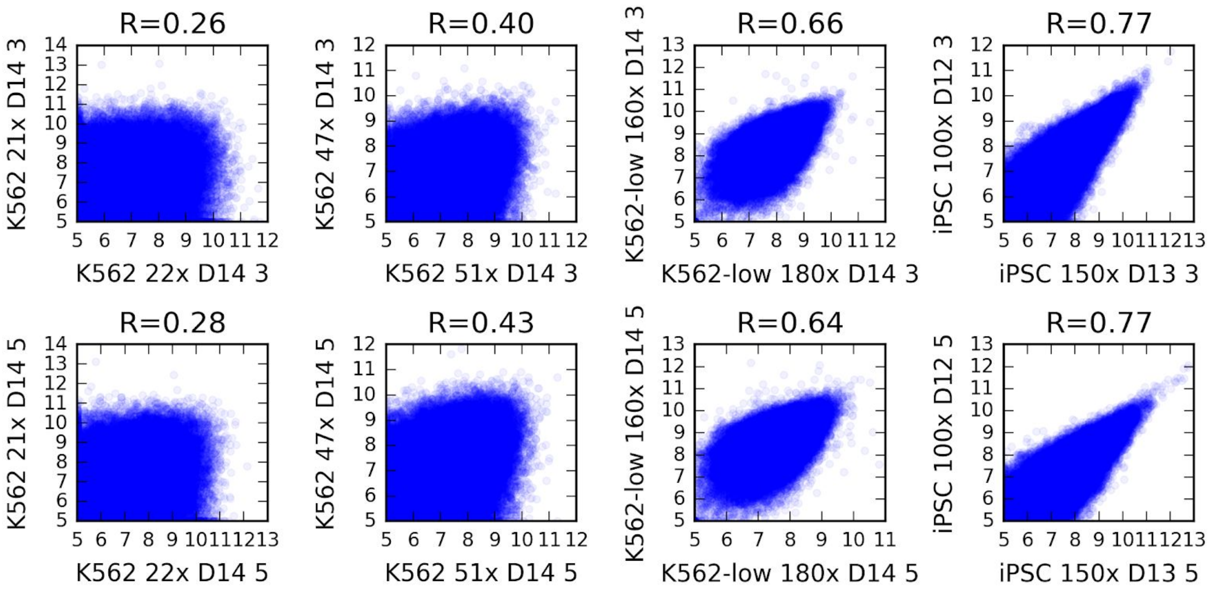
R=0.26
K562 21x D14 3
K562 22x D14 3
R=0.40
K562 47x D14 3
K562 51x D14 3
R=0.66
K562-low 160x D14 3
K562-low 180x D14 3
R=0.77
iPSC 100x D12 3
iPSC 150x D13 3
R=0.28
K562 21x D14 5
K562 22x D14 5
R=0.43
K562 47x D14 5
K562 51x D14 5
R=0.64
K562-low 160x D14 5
K562-low 180x D14 5
R=0.77
iPSC 100x D12 5
iPSC 150x D13 5

B.

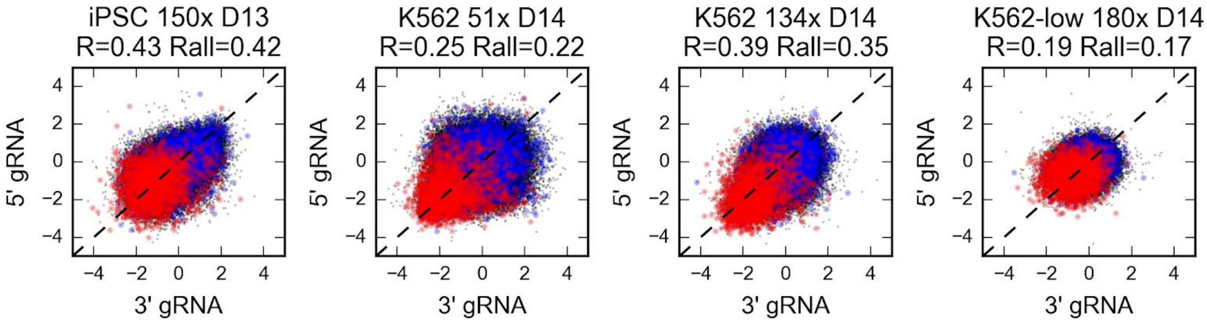
iPSC 150x D13
R=0.43 Rall=0.42
K562 51x D14
R=0.25 Rall=0.22
K562 134x D14
R=0.39 Rall=0.35
K562-low 180x D14
R=0.19 Rall=0.17
5' gRNA
3' gRNA

**C.**

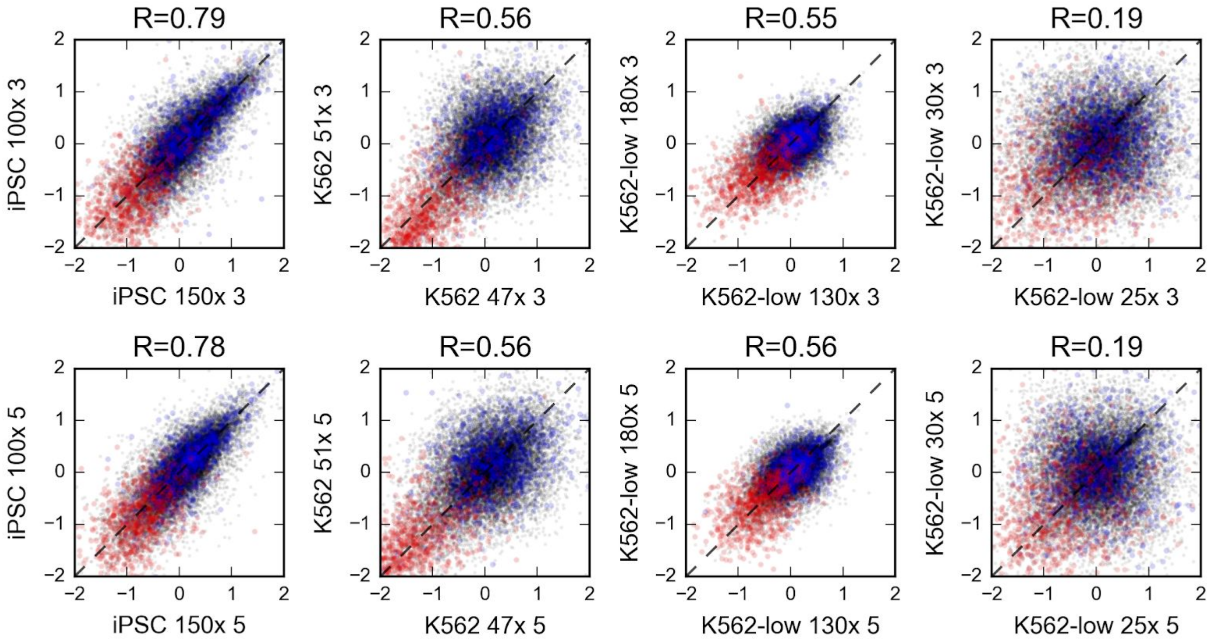


**D.**

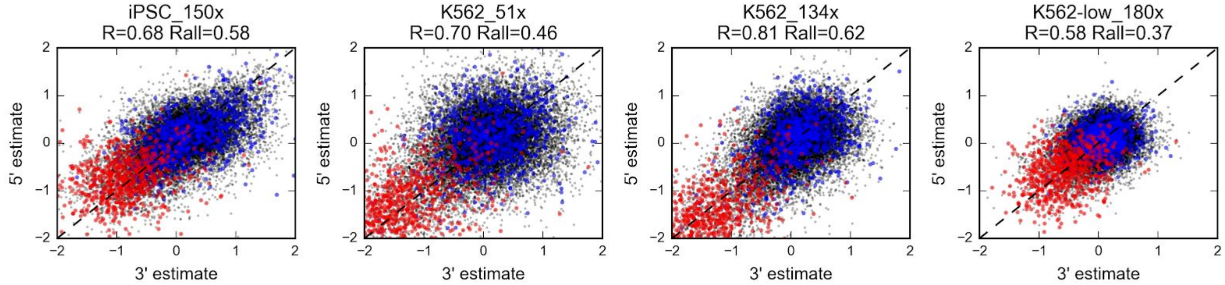


**Figure S5**. Screen results are reproducible for gRNA counts across replicates (A), gRNA effects across the two positions (B), gene effects across replicates (C), and gene effects across two positions (D). **A.** Log2-scale gRNA count in replicate 1 (x-axis) and replicate 2 (y-axis) for the 60,000 gRNAs in the 3' position (top row) and 5' position (bottom row) for four samples representing the range of coverages and cell types (columns). Sample names: cell type (K562, K562-low (low Cas9 efficacy), iPSC), coverage, screen timepoint. **B.** As in Figure 2A. Title: sample name as in panel A, Pearson's correlation of Hart essential and non-essential genes (R), and all genes (Rall). **C, D.** As A, B but for JACKS gene essentiality estimates.

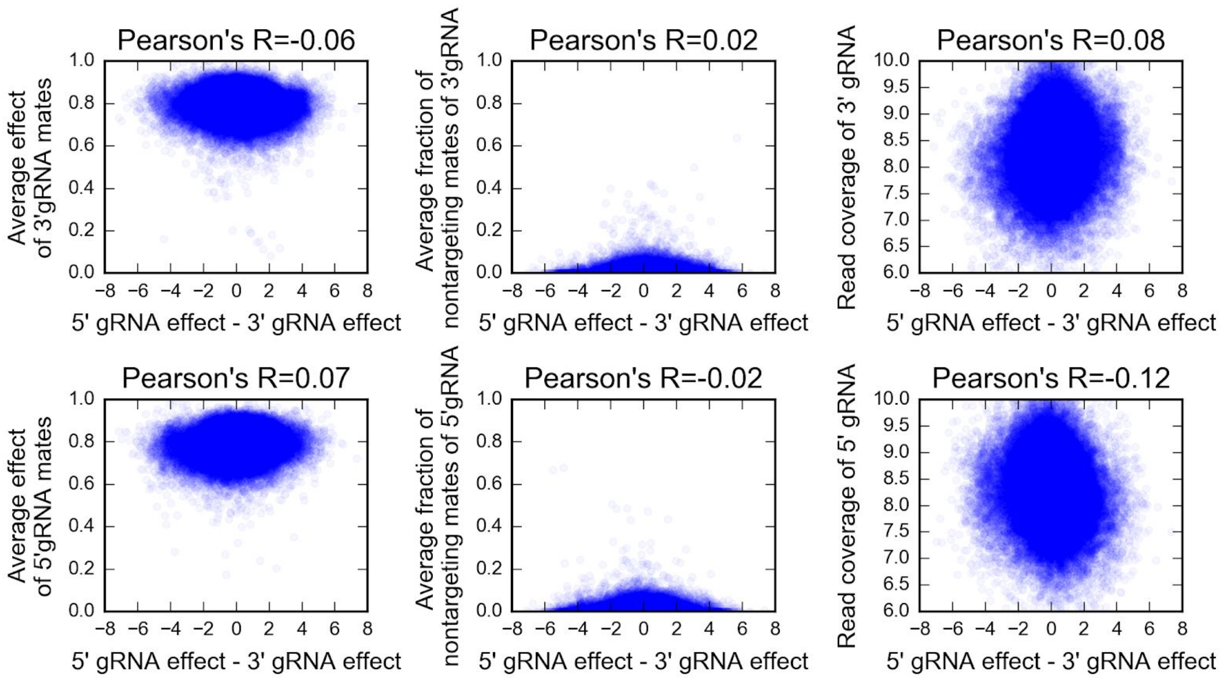


**Figure S6**. Differences in gRNA log-fold change compared to plasmid control between 5’ and 3’ gRNAs in iPSCs (150x screen, day 13, x-axis) are not explained by pairing to detrimental gRNAs (y-axis: average linear scale fitness effect of pair mates, left column), pairing to beneficial gRNAs (y-axis: average fraction of non-targeting gRNAs, middle column), or low coverage (y-axis: log2-scale read coverage, right column) for 3’ gRNA (top row) or 5’ gRNA (bottom row).

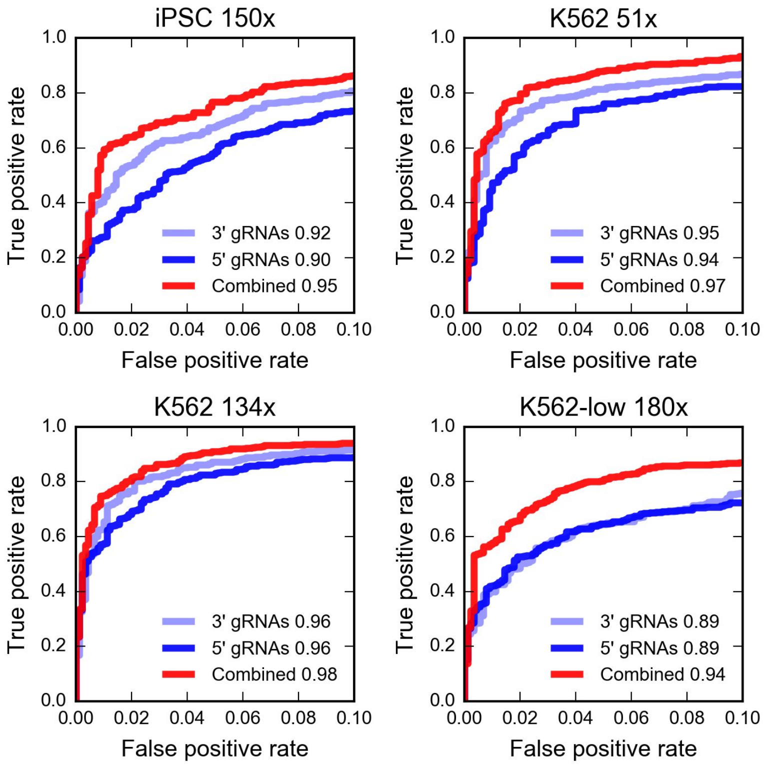


**Figure S7**. Performance improvement by combining information from both gRNA positions. True positive rate (y-axis) at increasing false positive rate (x-axis; zoomed to range 0 to 0.1) when using JACKS gene essentiality estimates to classify Hart gold standard essential genes from non-essential ones using only 3’ gRNAs (light blue), only 5’ gRNAs (dark blue), or a combination of both (red). Panels: representative samples from different cell lines. Area under the curve denoted in legend.

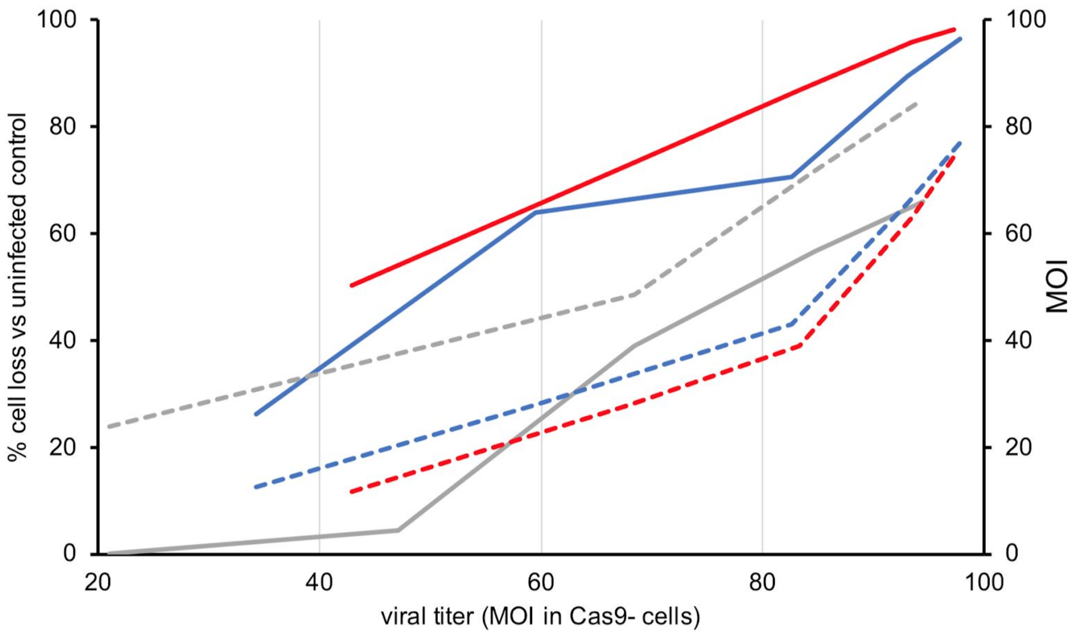


**Figure S8**. Additional double strand breaks do not lead to substantial additional cell loss. We performed cell metabolic activity assays to compare the effect on fitness of single- versus double-gRNA libraries. Percent cell loss compared to uninfected control (solid lines, left y-axis) and percent MOI (dashed lines, right y-axis) for different viral titer of Cas9-negative iPSCs (MOI, x-axis), for a library composed of non-targeting gRNAS (gray), single gRNA Yusa 1.0 (blue) and the double gRNA minimized library (red). Five days post-infection iPSC-Cas9 cells, infected at MOI of 0.28 (typical of genome-wide screens) showed an overall reduction in viability, with a slightly larger effect for the minimized library (27±5% surviving cells vs. uninfected) compared to Yusa 1.0 (36±10%). Further, viral amounts producing a MOI = 0.28 for both minimized and Yusa 1.0 libraries, displayed respectively MOI = 0.68 and 0.59 in iPSC-wild type cells, give an estimate of 25% larger cell loss upon infection for the double gRNA library compared to a single gRNA one.

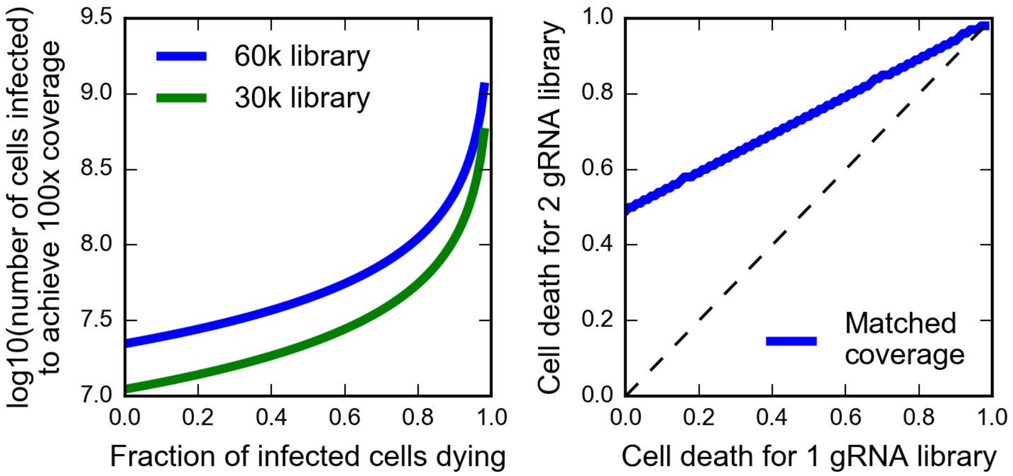


**Figure S9**. An effectively smaller library reduces scale only if it does not induce too much additional cell death. Left panel: log10-scale number of cells required for input into infection (y-axis) to achieve 100x coverage of each gRNA given the fraction of infected cells that die (x-axis) for a 60,000 gRNA library (blue) and 30,000 gRNA library (green), assuming 10% of cells die upon transduction from non-specific causes, and MOI of 0.3. The formula is *cells = library_size*coverage / (MOI*(1 - infected_cell_death)*(1-nonspecific_infection_death)),* where the numerator is the ultimate number of required cells, and the denominator accounts for the various types of cell loss. Right panel: “isocoverage” line (blue) at which two-gRNA and one-gRNA library have the same effective coverage using the same number of cells, for different cell death fractions upon infection for single gRNA library (x-axis) and double gRNA library (y-axis).

**A**

5’ gRNA oligonucleotide pool

GACGTCCAGAGCACAGATGGNNNNNNNNgatatcGCTTTATATATCTTGTGGAAAGGACGAAACAC
C**GNNNNNNNNNNNNNNNNNNN**GTTTGGgtcttcTCCGATGGTCTCCCTCGACAAACTCGTCTGAGC
TGTCATGGTCG

**B**

3’ gRNA oligonucleotide pool

GACGTCCAGAGCACAGATGGNNNNNNNNATCGATCTCCGATGGTCTCCCTCGACAAACTgaagacA
CTTTG**GNNNNNNNNNNNNNNNNNNN**GTTTAAGAGCTATGCTGGAAACAGCAgatatcCGTCTGAGC
TGTCATGGTCG

**C**

Scaffold and mU6 promoter, G-block

TATGAGGACGAATCTCCCGCTTATGAAGACCTGTTTAAGAGCTATGCTGGAAACAGCATAGCAAGT
TTAAATAAGGCTAGTCCGTTATCAACTTGAAAAAGTGGCACCGAGTCGGTGCTTTTTTTGTACTGA
GTCGCCCATCTAGAGATCCGACGCCGCCATCTCTAGGCCCGCGCCGGCCCCCTCGCACAGACTTGT
GGGAGAAGCTCGGCTACTCCCCTGCCCCGGTTAATTTGCATATAATATTTCCTAGTAACTATAGAG
GCTTAATGTGCGATAAAAGACAGATAATCTGTTCTTTTTAATACTAGCTACATTTTACATGATAGG
CTTGGATTTCTATAAGAGATACAAATACTAAATTATTATTTTAAAAAACAGCACAAAAGGAAACTC
ACCCTAACTGTAAAGTAATTGTGTGTTTTGAGACTATAAATATGCATGCGAGAAAAGCCTTGTTTG
AGGTCTTCTACAAGCACACGTTTGTCAAGACC

**Figure S10**. Sequence of gRNA oligonucleotide pools and G-block encoding scaffold and mU6 promoter. Primer binding sequences used for amplification of the oligonucleotide pools (A, B) or the G-block (C) are in black, underlined. $[N]_8$ regions in the oligonucleotide pools have different sequences to allow for the specific amplification of subpools. Promoter regions are in green, underlined. Scaffold sequences are in blue, underlined. Common sequence for fusion PCR is in red, underlined. gRNAs $G[N]_{19}$ are in bold. EcoRV and BbsI restriction sites are in lowercase. See Table S5 for details of primer sequences.

**Table S1**. gRNA sub-pools in the minimized genome-wide library

| gRNA sub-pool | Count |
|---|---|
| gRNA #1, essential genes | 5546 |
| gRNA #2, essential genes | 5544 |
| gRNA #3, essential genes | 5496 |
| gRNA #1, non-essential genes | 13713 |
| gRNA #2, non-essential genes | 13707 |
| gRNA #3, non-essential genes | 13610 |
| gRNA #4, iPSC-specific non-essential genes | 1986 |
| Non-targeting controls | 398 |

**Table S2.** Quality metrics for the performed screens analysed with alternative approaches. The first three columns describe the screen and the analysis method. First column - Sample identifier (cell line, coverage, timepoint, optionally 3' or 5' gRNA set used). Second column - encoding of the 3' and 5' position gRNAs. "Yusa" - Yusa 1.0 library. "grnas" - the G = about 60,000 different 3' and 5' gRNAs are treated as independent, resulting in a 2*G = ~120,000-long read count file for the N screens. "Samples" - the G 3' and 5' gRNAs are treated as different samples, resulting in a G = ~60,000-long read count file for 2*N screen samples. "samples_total" - average score of the 5' and 3' samples for the same screen. Third column - number of gRNAs used in gene effect calculation. 1 - single gRNA. 2- two gRNAs. 4 - all gRNAs (3 for most genes, additional one for ones highly expressed in iPSCs (Methods)). "gRNA LFC" - metrics calculated from gRNA log-fold changes only. "Gene LFC" - metrics calculated with average gRNA log-fold change as essentiality estimate, rather than JACKS score. Fourth column - Delta AUC. Columns 5-8: 1-recall at different false positive rates (0.05 to 0.5). Columns 9-11: partial area under the curve (pAUC) at different false positive rate cutoffs (0.1, 0.2, 1.0). Columns 12-14: false positive rate at different true positive rates (0.75, 0.80, 0.85).

**Table S3.** List of pathways and processes enriched for 250 top genes ordered by differential essentiality between K562 and iPSC, with the genes more essential in K562.

**Table S4.** As table S3, but for genes more essential in iPSCs.

**Table S5. Primer sequences**

| Primer | Sequence | Application |
|---|---|---|
| #221 | GACGTCCAGAGCACAGATGG | Library cloning |
| #270 | CGACCATGACAGCTCAGACG | Library cloning |
| #526 | AGTTTGTCGAGGGAGACCATCGGAG | Library cloning |
| #527 | CTCCGATGGTCTCCCTCGACAAACT | Library cloning |
| #545 | TATGAGGACGAATCTCCCGCTTATG | gBlock amplification |
| #546 | GGTCTTGACAAACGTGTGCTTGTAG | gBlock amplification |
| #1 | ACACTCTTTCCCTACACGACGCTCTTCCGATCTCTTGTGGAAAGGACGAAACA | Sequencing, gRNA library amplification |
| #2 | TCGGCATTCCTGCTGAACCGCTCTTCCGATCTCTAAAGCGCATGCTCCAGAC | Sequencing, gRNA library amplification |
| #430 | TCGGCATTCCTGCTGAACCGCTCTTCCGATCTGCTGTTTCCAGCATAGCTCTTAAAC | Sequencing, gRNA library amplification |
| #463 | TCGGCATTCCTGCTGAACCGCTCTTCCGATCTGCACCGACTCGGTGCCACTT | Sequencing, gRNA library amplification |
| #605 | ACACTCTTTCCCTACACGACGCTCTTCCGATCTATGCATGCGAGAAAAGCCTTGTTTG | Sequencing, gRNA library amplification |
| #15 | AATGATACGGCGACCACCGAGATCTACACTCTTTCCCTACACGACGCTCTTCCGATCT | Sequencing, indexing PCR |
| indexing | CAAGCAGAAGACGGCATACGAGAT$N_{11}$GAGATCGGTCTCGGCATTCCTGCTGAACCGCTCTTCCGATCT | Sequencing, indexing PCR |
| #16 | TCTTCCGATCTCTTGTGGAAAGGACGAAACACCG | Sequencing |
| #436 | TCGGCATTCCTGCTGAACCGCTCTTCCGATCT | Sequencing |
| #619 | TCTTCCGATCtATGCATGCGAGAAAAGCCTTGTTTGG | Sequencing |
